# Isotype specific loss of HP1α but not of HP1β uncovers genomic regions that behave as HP1α-dependent common fragile sites

**DOI:** 10.64898/2026.08.14.744815

**Authors:** Karen Yaacoub, Thanh Nhan Nguyen, Eric Julien, Florence Cammas

## Abstract

HP1 proteins are highly evolutionarily conserved chromatin-associated factors known to play essential roles in genome stability and nuclear organization. In mammals, three HP1 isoforms, HP1α, HP1β and HP1γ, have been described, but their individual functions remain incompletely characterized. Here, we inactivated HP1α or HP1β in different cell lines and quantified chromosomal breaks on metaphase spreads in the presence or absence of aphidicolin-induced replication stress. Loss of HP1α, but not of HP1β, led to a significant increase of chromosomal breaks on chromosome arms and within pericentromeric heterochromatin under these conditions. Mechanistically, loss of HP1α was associated with a reduction in replication fork velocity, suggesting that HP1α deficiency induces a replication stress that sensitizes specific genomic loci to replication perturbation. Consistent with this, HP1α loss was associated with a moderate but consistent increase in γH2AX and 53BP1 foci, an increased occurrence of DNA synthesis during mitosis, and enhanced recruitment of FANCD2, all recognized as hallmarks of common fragile site (CFS) expression. In addition, rescue experiments using a chromodomain mutant HP1α (V22M) unable to bind H3K9me3 indicated that HP1α protective function over these specific foci did not require its interaction with this histone mark. Altogether, these data indicate that, independently of its binding to H3K9me3, HP1α stabilizes specific genomic regions that behave as HP1α-dependent fragile sites, at least in part by regulating replication fork progression, limiting mitotic DNA synthesis possibly by competing with FANCD2 for chromatin access at these regions.

## Introduction

Genome stability is essential for the proper functioning of cells, tissues, and organisms and is associated with chromatin and nuclear organization (1). Among the chromatin-associated factors implicated in genome stability, HP1 proteins have emerged as key regulators of nuclear organization and genomic integrity. These proteins are highly conserved throughout evolution, from yeast to mammals, and are involved in all nuclear processes involving chromatin organization throughout heterochromatin with increasing evidence that they are also essential within euchromatin. The pleiotropic functions of HP1s rely on their structural organization, with a N-terminal Chromodomain (CD) allowing these proteins to associate with chromatin via recognition of the repressive Histone H3 lysine 9 tri-methylated (H3K9me3) mark, and a C-terminal ChromoShadow domain (CSD) involved in HP1 dimerization, providing a platform for the recruitment of multiple proteins to chromatin (2). Beside this canonical binding of HP1 to H3K9me3, HP1 also bind to genomic loci independently of H3K9me3 and this binding is often involved in regulation of gene expression (3). In mammals, there are three HP1 isoforms— HP1α, β, and γ. Despite sharing a conserved structural organization, early studies established that the three isoforms occupy distinct nuclear compartments, HP1α being predominantly heterochromatic, HP1β distributed between eu- and heterochromatin and HP1γ being mostly euchromatin, suggesting that their functional specificities are at least partly determined by their chromatin environment (4). Accordingly, the three mammalian HP1 isoforms exhibit specific non redundant physiological functions with HP1α being essential for the maintenance of T cell identity (5,6), HP1β being required for development of the cerebral neocortex and neuromuscular junctions (7) and HP1γ essential for spermatogenesis (8). At the molecular level, HP1α, HP1β and HP1γ play roles in a wide range of nuclear processes including transcriptional regulation, DNA replication and DNA repair, yet their individual contributions as well as their nuclear organization toward these processes remain incompletely understood (9). In line with HP1 functions in all chromatin associated processes, alterations of HP1 expression are increasingly found to be associated cancer development (10,11).

More recently, it has become clear that HP1α and HP1β direct distinct DNA repair pathway choices in response to double-strand breaks occurring within heterochromatin, further highlighting the non-redundant nature of their functions (12). Together, these findings underscore the importance of studying each HP1 isoform individually, and raise the question of whether their functional specificities extend to the protection of specific genomic regions in stressed conditions. In this regard, although eukaryotic cells evolved complex protein networks dedicated to genome protection, there are specific regions that are naturally unstable. Among them, the so-called Common Fragile Sites (CFSs) are regions that, upon replication perturbation, behave as hotspots for chromosomal rearrangements such as deletions, translocations and amplifications that are recurrent in many cancers (13). Interestingly, some genomic features underlying CFS closely resemble those characterizing HP1-bound regions including that both are generally associated with very large genes enriched in AT-dinucleotide sequences and both exhibit delayed replication (13–15). Here, based on these similarities, we investigated whether HP1α and/or HP1β isoforms contribute to the protection of specific genomic loci against replication stress-induced instability, using loss-of-function approaches combined with aphidicolin-induced replication stress to unmask potential fragility.

## Results

### Loss of HP1α but not of HP1β leads to increase chromosomal breakage in response to aphidicolin-induced replication stress

Because HP1 binding sites share several features with common fragile sites (CFS) (16), we investigated whether loss of HP1 could promote chromosomal breaks under replicative stress. To this end, we inactivated the genes encoding HP1α or HP1β (*cbx5* and *cbx1*, respectively) using the genome editing system CRISPR–Cas9 in the mouse hepatocyte AML12 cell line. Loss of HP1α (HP1αKO) and HP1β (HP1βKO) was verified by WB in the two respective sub- cloned cell lines (Figure 1A). Loss of either HP1α or HP1β in AML12 did not induce any significant alteration of the proliferative properties of these cells (Figure S1A). For all further experiments we treated cells with a moderate dose (1.2 µM) of APH for 20h. This condition was determined empirically as the lowest dose of aphidicolin promoting a significant increase of chromosomal breakage in the parental AML12 Wild-type (WT) cell line that we used in this study. Metaphase spreads were subsequently prepared, stained with DAPI (4′,6-diamidino-2- phenylindole dihydrochloride), and analyzed for chromosomal breaks by fluorescent microscopy which allowed us to separate breaks on chromosomal arms (green arrows on Figure 1B) and breaks within pericentromeric heterochromatin (PHC, red arrows on Figure 1B) nearby chromocenters that are characterized as DAPI dense regions (shown as white arrows on Figure 1B). In this condition, loss of HP1α (HP1αKO) led to a significant increase of the number of breaks on chromosomal arms as compared to WT cells, whereas the loss of HP1β (HP1βKO) had no significant effect with even a tendency of a decrease number of breaks as compared to WT cells (Figure 1E). In agreement with the known role of HP1 for heterochromatin maintenance, the number of chromosomal breaks at PCH was higher in both HP1αKO and HP1βKO cell lines as compared to WT cells, although the difference did not reach statistical significance in the case of the HP1βKO cell line (Figure 1F). To ensure that the observed phenotype was specifically associated with the loss of HP1α and not to off-target effects of the CRISPR–Cas9 strategy, we stably introduced a transgene allowing doxycycline (Dox)-inducible expression of Flag-tagged HP1α (f-HP1α) protein into HP1αKO AML12 cells (HP1αKO-f-WT). Immunofluorescence (IF) analysis revealed that upon treatment of HP1αKO- f-WT cells with doxycycline, f-HP1α was highly expressed and preferentially localized in DAPI-dense foci, the hallmark of PCH in mouse cells (Figure 1C, upper panels). WB analysis showed that f-HP1α is expressed at a much higher level than the endogenous protein and that some low level of f-HP1α expression could be seen even in absence of Dox treatment (Figure 1D). This low level of f-HP1α expression appears to be sufficient to decrease the number of chromosomal breaks within arms even below the number of chromosomal breaks observed in WT cells (Figure 1E). This effect was not significantly changed by further treatment with Dox (Figure 1E). In contrast, the number of chromosomal breaks at PCH was not rescued by re-expression of f-HP1α neither in presence or absence of Dox (Figure 1F).

**Figure 1.**
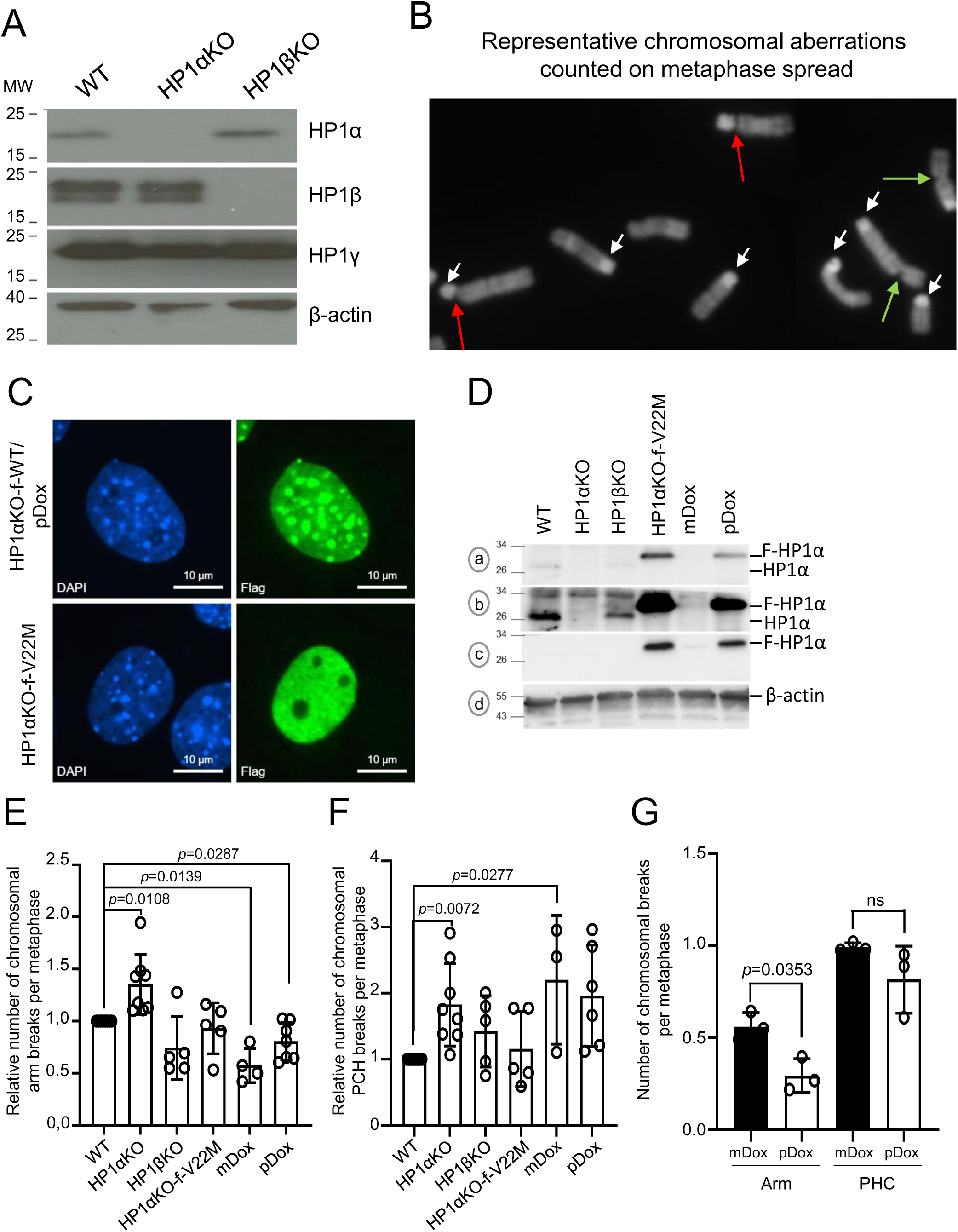
: HP1α is involved in the stabilization of specific genomic loci in response to aphidicolin-induced replication stress. (A) Western Blot (WB) analysis of WCE from mouse hepatocyte AML12 WT and CRISPR-Cas9 derived and sub-cloned HP1αKO and HP1βKO cell lines with the indicated antibodies. The molecular weights (MW) are shown on the left of the blots (B) Representative image of the chromosomal abnormalities counted in this analysis with breaks throughout chromosomal arms (green arrows) and breaks around pericentromeric heterochromatin (PCH, red arrows) on a metaphasic plate. The DAPI-dense centromeric regions are shown with white arrows. (C) IF analysis of HP1αKO AML12 cell line re-expressing in a doxycycline inducible manner a flag-tagged HP1αWT (HP1αKO-f-αWT) in presence of 1mg/ml doxycycline (pDox) allowing the transient expression of flag-tagged HP1α or the stably established cell line expressing flag-tagged HP1αV22M (HP1αKO-f-V22M) with an anti-Flag antibody (green). Nuclei are stained with DAPI (blue). (D) WB analysis of AML12 WT, HP1αKO, HP1βKO, HP1αKO-f-V22M and HP1αKO-f-WT in presence (pDox) or absence (mDox) of 1mg/ml doxycycline with anti-HP1α (blot a for short exposure and blot b for long exposure), anti-Flag (blot c) and anti-β-actin (blot d) antibodies. (E) Quantification of the breaks within chromosomal arms of AML12 WT, HP1αKO, HP1βKO, HP1αKO cell line re-expressing the flag-tagged HP1αV22M (HP1αKO-f-V22M) or f-HP1α (HP1αKO-f-WT) cell line in presence (pDox) or absence (mDox) of 1mg/ml doxycycline and treated for 20h with 1.2µM APH. (F) Quantification of breaks around PCH for the same cell lines and in the same conditions as (E). For panels E and F, the mean values for each condition were normalized to WT in each independent experiment (n=8). An average of around 50 metaphases were counted per condition. Statistical significance was assessed using a one-sample t-test against the theoretical value of 1, the *p-*values correspond to comparison between WT and the different cell lines and were calculated using GraphPad Prism software. (G) Counting of the number of breaks within chromosomal arms and PHC on metaphasic plates of HP1αKO-f-WT AML12 cells treated for 20h with 1.2µM APH in presence (white bars, pDox) or absence (black bars, mDox) of 1mg/ml doxycycline in a medium deprived of residual tetracycline. The mean values for three independent experiments are shown on the graph. One-way ANOVA statistical test was used and ns correspond to *p*value>0.05. An average of around 50 metaphases were counted per condition.

For this reason, we performed a specific set of experiments measuring the number of chromosomal breaks comparing Dox treated and non-treated HP1αKO-f-WT cells in a medium deprived of residual tetracyclin. Under APH treatment, induction of f-HP1α expression in HP1αKO-f-WT cells led to a significant reduction of the number of chromosomal breaks as compared with APH-treated cells in absence of Dox (Figure 1G, compare pDox with mDox), indicating that the increased number of chromosomal breaks observed in HP1αKO cells was indeed a consequence of the loss of HP1α. Although in these conditions we also observed a decrease of the number of breaks at PCH, this difference did not reach statistical significance, suggesting that once established, breaks induced by loss of HP1α within PHC cannot be efficiently repaired by re-expression of f-HP1α (Figure 1F and G).

To gain insight into the mechanisms by which HP1α contributes to prevent chromosomal breakage in response to APH-induced replicative stress, we generated AML12 HP1αKO cells stably expressing a flag-tagged HP1α protein mutated in its chromodomain-mutant (V22M), a mutation known to abolish HP1α binding to H3K9me3 (17). IF analysis revealed that, as expected and in contrast to WT f-HP1α, f-HP1αV22M failed to localize to the DAPI-dense foci (Figure 1C, lower panel). WB analysis showed that f-HP1α (pDox) and f-HP1αV22M are expressed at similar levels and higher levels than the endogenous HP1α protein (figure 1D). Expression of f-HP1αV22M in HP1αKO cells treated with APH led to a significant reduction of the number of chromosomal breaks within arms to reach levels comparable to those observed in WT AML12 cells (Figure 1E). Furthermore, in contrast to f-HP1α, f-HP1αV22M expression led to a decreased number of chromosomal breaks at PCH to a level similar to the one observed in WT cells (Figure 1F).

To assess whether this protective role of HP1α against chromosomal breaks induced by replicative stress extends beyond a single cell line, we used our previously described *cbx5* inactivated human hepatocellular carcinoma HepG2 cell line (18). We further performed CRISPR-Cas9 inactivation of *cbx1* in this same HepG2 parental WT cell line. After validation of the inactivation of *cbx5* (HP1αKO) or *cbx1* (HP1βKO) by WB analysis (Figure S1B), we analyzed the number of breaks on metaphases as described above. As observed in AML12 cells, loss of HP1α led to an increase number of chromosomal breaks in APH treated cells as compared to WT cells although this increase did not reach statistical significance (Figure S1C). As in AML12 cells, loss of HP1β led to a trend to decrease the number of breaks as compared to WT cells (Figure S1C). Similarly, Mouse Embryonic Fibroblasts (MEFs) derived from HP1αKO embryos exhibited increased chromosomal breakage upon APH exposure compared to control *HP1α^+/-^* MEF (Figure S1D-F), although, in contrast to AML12 cells, loss of HP1α in MEF cells led to a significant decrease of cell proliferation (Figure S1H).

Altogether, these data demonstrated that, under APH-induced replication stress, loss of HP1α, but not of HP1β, increased the number of breaks within chromosomal arms on metaphase spreads whereas the loss of either HP1α or HP1β led to increased number of chromosomal breaks within PCH. This finding supports a specific role for HP1α in safeguarding discrete genomic regions that behave as HP1α-dependent fragile sites. Importantly, this protective function of HP1α is uncoupled from its canonical interaction with H3K9me3 and independent of the proliferative capacity of the cells.

### DNA Damage Response is moderately enhanced upon loss of HP1α

CFS are characterized by the activation of the response to Double Strand Breaks (DSB) via activation of the ATR pathway through phosphorylation of the kinase Chk1 (19). We therefore tested whether the activation of this pathway was altered upon loss of HP1α. WB analysis of Chk1 serine 345 phosphorylation showed that ATR/Chk1 pathway is activated in both WT and HP1αKO AML12 cells at a slightly higher level in this latter cell line upon APH treatment (Figure 2A). Consistent with this, IF analysis showed that the established marker of DNA breaks, γH2AX (phosphorylated histone H2AX on serine 139) is detectable in non-stressed WT, HP1αKO, and HP1βKO cells and increased upon APH treatment in all cell lines with a 1.2-fold increase in HP1αKO as compared to WT cells treated cells (Figure 2B-C). To further analyze the response of AML12 cells to APH treatment, we also tested the expression of 53BP1, whose foci are known to mark DSB repair but also largely correspond to incompletely replicated CFS loci in G1 phase (20). As for γH2AX, 53BP1 foci are detectable in non-stressed WT, HP1αKO, and HP1βKO cells and increased upon APH treatment in all cell lines with a 1.4-fold increase in HP1αKO as compared to WT treated cells (Figure 2B and D). These findings indicated that the DNA damage response pathways upon APH treatment remain functional in the absence of either HP1 proteins and suggest that the increased number of γH2AX and 53BP1 foci observed in HP1αKO cells, although not statistically significant, correspond to incompletely replicated regions behaving as CFS that are rare events throughout the genome.

**Figure 2.**
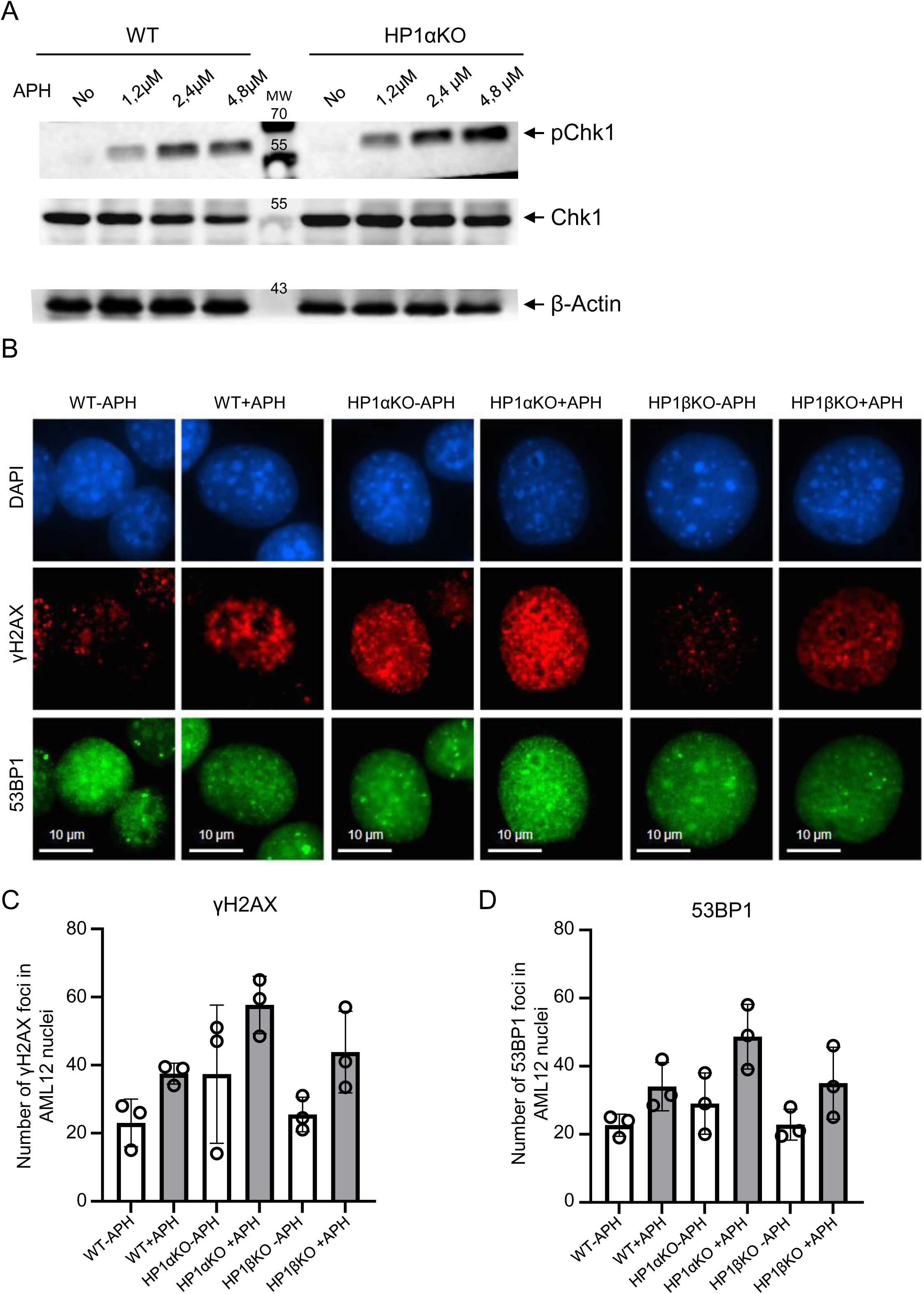
: Loss of HP1α leads to a moderate increase of the response to DNA damage. (A) Representative WB analysis of Chk1 serine 345 phosphorylation (pChk1) as a marker of ATR/Chk1 pathway activation and of the total amount of Chk1 (Chk1) in response to increasing doses of APH in WCE of WT and HP1αKO AML12 cells. The molecular weight (MW) are shown between the two cell lines (B) Representative IF analysis of phosphorylated histone H2AX on serine 139 (γH2AX) and 53BP1 in absence (-APH) or presence (+APH) of 1.2µM APH in WT, HP1αKO and HP1βKO AML12 cells. (C) Quantification of the number of γH2AX foci in 3 independent experiments counting around 50 nuclei per condition. (D) Quantification of the number of 53BP1 foci in 3 independent experiments counting around 50 nuclei per condition. 53BP1 and γH2AX foci per nucleus in WT, HP1αKO and HP1βKO cells were assessed using the cell profiler software. Ordinary one-way ANOVA test was conducted for pairwise comparisons of the mean number of foci for the 3 independent experiments using the GraphPad Prism software. None of the comparison was statistically significant (*p*value>0.05).

### Replication fork velocity is decreased upon loss of HP1α

Since HP1 proteins are known to be directly implicated in the regulation of DNA replication, we hypothesized that the increased chromosomal breakage observed under APH-induced replicative stress in HP1αKO cells could be associated with alterations of the DNA replication fork velocity. To investigate this hypothesis, we performed DNA fiber assays to measure replication fork progression in WT and HP1αKO AML12 cells. The average of the median length of replication tracks labeled with IdU and CldU were 1.3-fold shorter in HP1αKO cells compared to those measured in WT controls, indicating a reduction in replication fork speed upon HP1α loss (Figure 3A-B). This result demonstrates that HP1α deficiency impairs replication fork progression. DNA replication is tightly regulated by numerous factors, some of which have been reported to interact with HP1 proteins. To determine whether HP1α loss affects the expression or chromatin association of key replication regulators, we analyzed the levels and the chromatin association of ORC1, ORC2 (21), MCM10 (22), TRIM28 (23), and the other HP1 isoforms (HP1β and HP1γ) via cellular fractionation followed by WB analysis. No significant differences in expression or chromatin association of any of these proteins were detected between HP1αKO and WT cells (Figure S2). Collectively, our data demonstrate that HP1α deficiency induced a mild replication stress characterized by reduced replication fork velocity which is associated with an increase number of chromosomal breaks observed on metaphasic spreads. Importantly, this phenotype occurs without alteration of the expression or chromatin recruitment of several critical replication factors known to associate with HP1, suggesting that HP1α influences replication dynamics through mechanisms beyond direct regulation of the recruitment of these proteins to chromatin.

**Figure 3:**
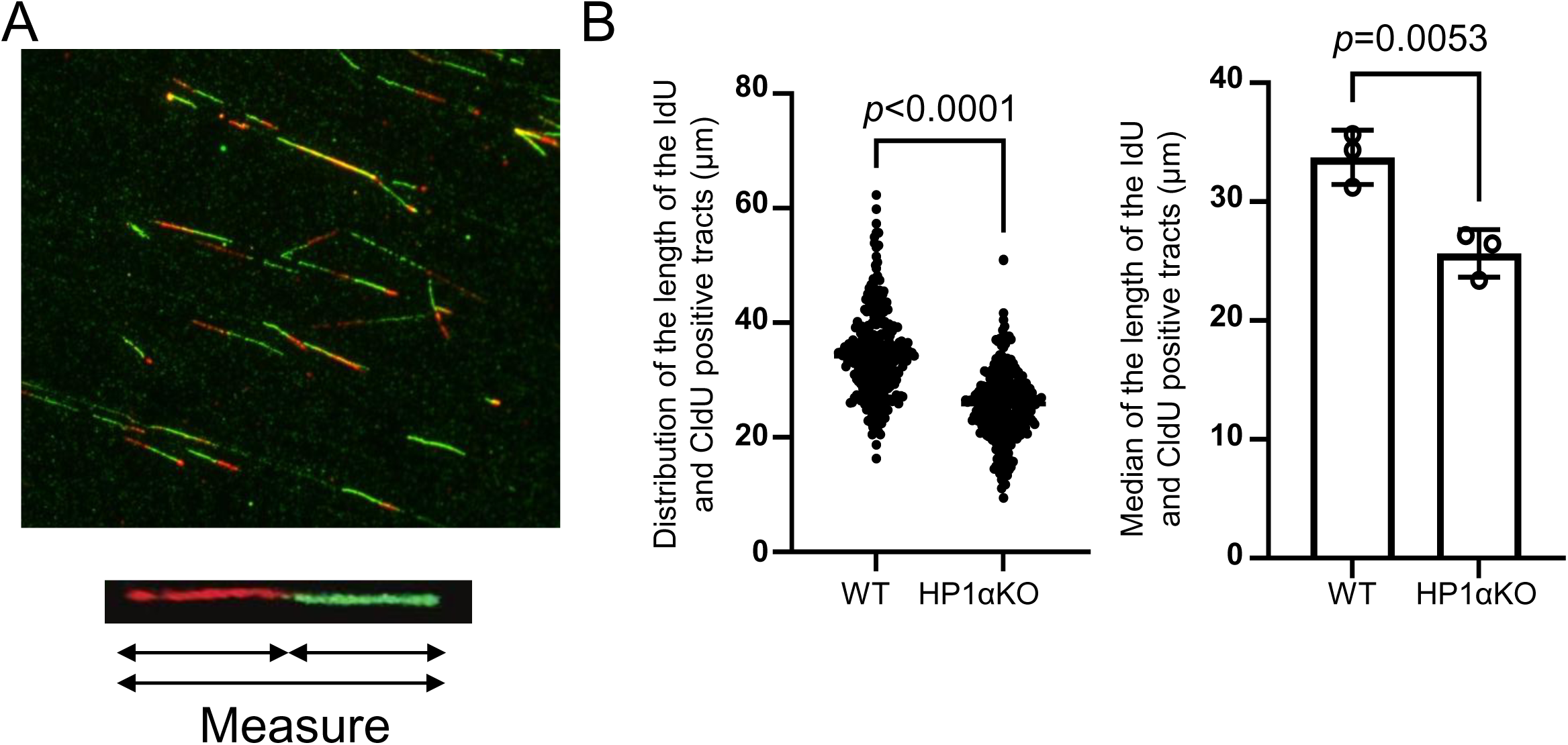
Replication fork velocity is impaired upon loss of HP1α in AML12 cells. WT or HP1αKO AML12 were grown in absence of treatment for 48h and then sequentially treated with IdU then with CldU (n=3). (A) Representative images of the IdU+CldU positive fibers that have been counted in this study (B) Graph representing the distribution of the size of the IdU+CldU positive fibers counted in the three independent experiments (left) and of the median size of the IdU+CldU positive fibers for the three independent experiments (right). Fibers were visualized and imaged by Zeiss Axio Imager M2 upright microscope (Zeiss) equipped with an Apochromat 40X objective (NA 1.4, immersion oil). About 100 tracts were counted in each experimental condition. The length of the tracks was analyzed using ImageJ software. Statistical analysis was performed with a paired t-test comparing the median length of the tracks for the 3 independent experiments using GraphPad Prism software.

### Loss of HP1α but not of HP1β leads to increase mitotic DNA replication in response to replicative stress

To maintain genome stability and ensure its accurate transmission, DNA replication must strictly be complete once and only once prior the beginning of mitosis. Upon mild replicative stress, DNA synthesis at late replication sites can overlap with mitotic chromosome condensation generating the phenomenon of mitotic DNA synthesis, called MiDAS (24). Common fragile sites (CFS) were shown to co-localize to sites of DNA synthesis during mitosis (25), an observation that was confirmed by genome-wide analyses showing that CFS are enriched in the MiDAS fraction under replicative stress conditions (26). Based on this literature and our observation that the loss of HP1α leads to decreased replication fork velocity, we hypothesized that the increased genome instability observed in HP1αKO cells under APH-induced replicative stress could result from an increased probability of these cells to complete replication during mitosis. To test this hypothesis, APH-treated WT, HP1αKO, and HP1βKO AML12 cells were synchronized at the G2/M transition and released in presence of the DNA synthesis marker 5-ethynyl-2’-deoxyuridine (EdU). EdU incorporation was revealed by click-it reaction and mitotic cells were detected with an anti-phosphorylated histone H3-S10 antibody (Figure 4A). As expected, based on the current literature, some AML12 WT mitotic cells displayed incorporation of EdU in these conditions (Figure 4A-B, EdU bright spots that are circled in red on the “detected” images, see materials and methods). This category of mitotic cells having incorporated EdU was specifically enriched in the HP1αKO cell line (1.87-fold-change) but not in HP1βKO cell line which behaved very similarly to WT (0.96-fold-change) (Figure 4A-B). To confirm that this elevated EdU incorporation was specific to mitosis, EdU incorporation during interphase was also assessed. Unlike mitotic cells, EdU incorporation was reduced in both HP1αKO and HP1βKO interphase cells compared to WT corroborating the decreased fork velocity observed in HP1αKO cells (Figure S3). Similar results were obtained in the human HepG2 cells with a specific increase of MiDAS activity in HP1αKO but not HP1βKO cells (Figure S4). These data thus indicated that the specific loss of HP1α but not of HP1β is associated with an increased DNA synthesis during mitosis in response to APH-induced replicative stress.

**Figure 4:**
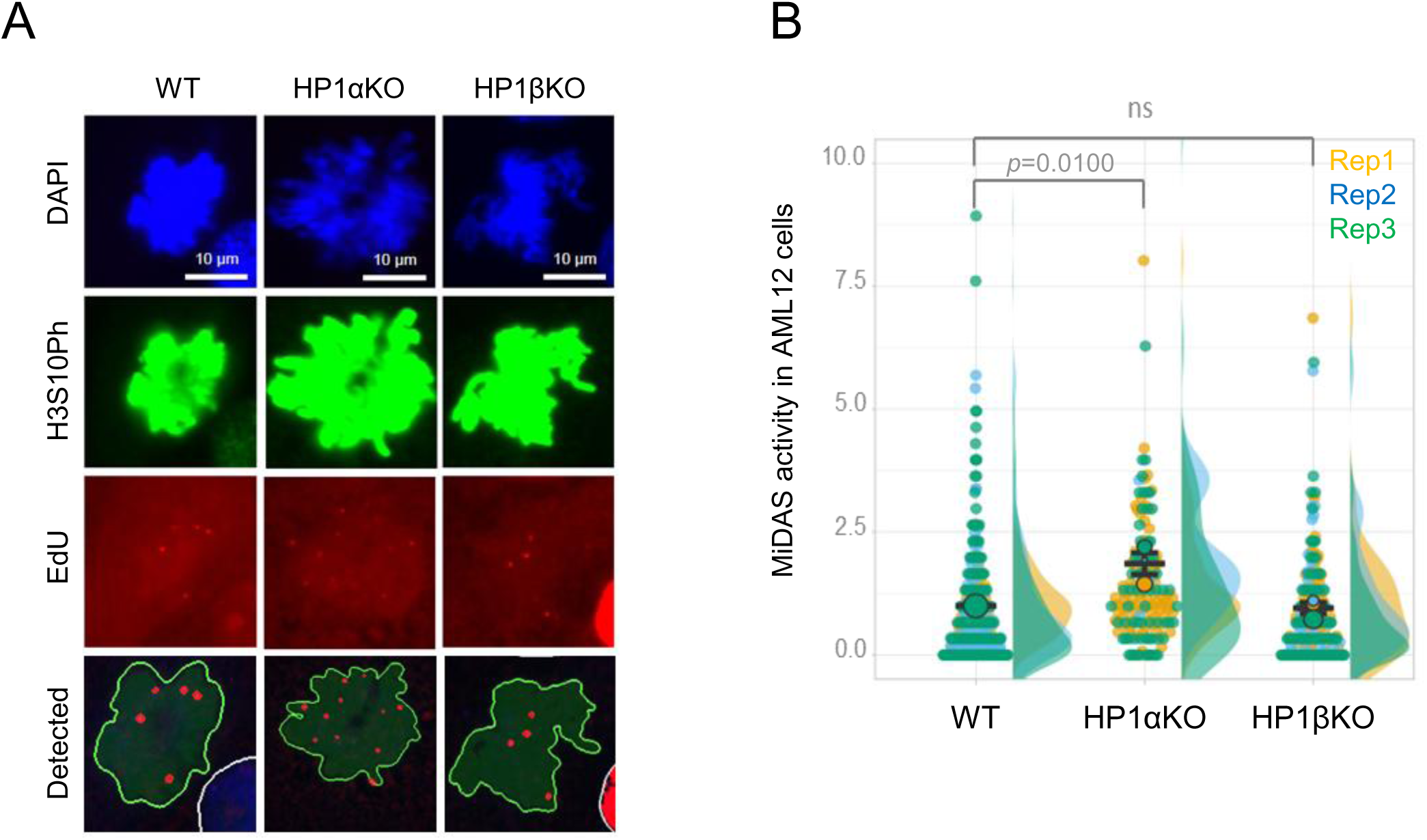
Loss of HP1α but not of HP1β leads to increase mitotic DNA replication in response to S-phase replicative stress. (A) representative microscopic images of WT, HP1αKO and HP1βKO AML12 nuclei (DAPI) after staining with anti-phospho H3S10 as a marker of mitotic cells (green), anti-EdU as a marker of DNA replication (red) antibodies. The red circles in the “detected” images correspond to the EdU spots quantified by cell profiler analysis in the mitotic cells (H3S10P positive nuclei outlined in green). (B) Superplot showing the distribution of the number of EdU foci per WT, HP1αKO and HP1βKO AML12 mitotic nucleus as well as the mean number of EdU foci per mitotic nucleus normalized to WT for the three independent experiments (Rep1 in yellow, Rep2 in blue and Rep3 in green). Statistical analysis shown on the graph has been performed as paired t-test on the normalized means to WT of three independent experiments. (rep1: WT, n=31, HP1αKO, n=82, HP1βKO, n=55; rep2 : WT, n=65, HP1αKO, n=4, HP1βKO, n=35 and Rep3 WT, n=212, HP1αKO, n=68, HP1βKO, n=122).

### Impact of loss of HP1α or HP1β on FANCD2 recruitment at MiDAS foci

Fanconi anemia group D2 protein (FANCD2) has been demonstrated to play an essential role in the regulation of MiDAS and to be specifically enriched at CFS loci upon replication stress (26). To test whether the recruitment of this protein on MiDAS foci was affected by the loss of HP1, we performed the same experiments as described above but adding a step of IF to detect FANCD2. As shown on figure 5A and B, FANCD2 is expressed at an equivalent level in all WT, HP1αKO and HP1βKO AML12 cells. Quantification of the number of FANCD2 foci within mitotic cells indicated that, as the number of MiDAS foci, FANCD2 foci were specifically increased in HP1αKO (79 mean number of foci/mitosis, two-fold change) but not in HP1βKO cells (35 mean number of foci/mitosis), as compared to WT cells (39 mean number of foci/mitosis) (Figure 5C). Further analysis specifically on MiDAS foci revealed that FANCD2 relative intensity to WT was higher in HP1αKO MiDAS foci (1.17-fold-change) and but not HP1βKO MiDAS foci which displayed an even lower intensity (0.88-fold-change) (Figure 5D). Because these data were obtained on two independent experiments, we did not perform statistical analysis, however considering the large number of nuclei analyzed, we believe that these differences are of biological significance. This experiment was also performed in human HepG2 cells (Figure S4). In contrast to AML12, we did not observe an increase number of FANCD2 foci within mitotic HP1αKO HepG2 nuclei, but rather a tendency of a decrease number of these foci. However, as in AML12, FANCD2 intensity was increased in HepG2 HP1αKO and to a lesser extent in HP1βKO MiDAS foci as compared to WT (Figure S4).

**Figure 5:**
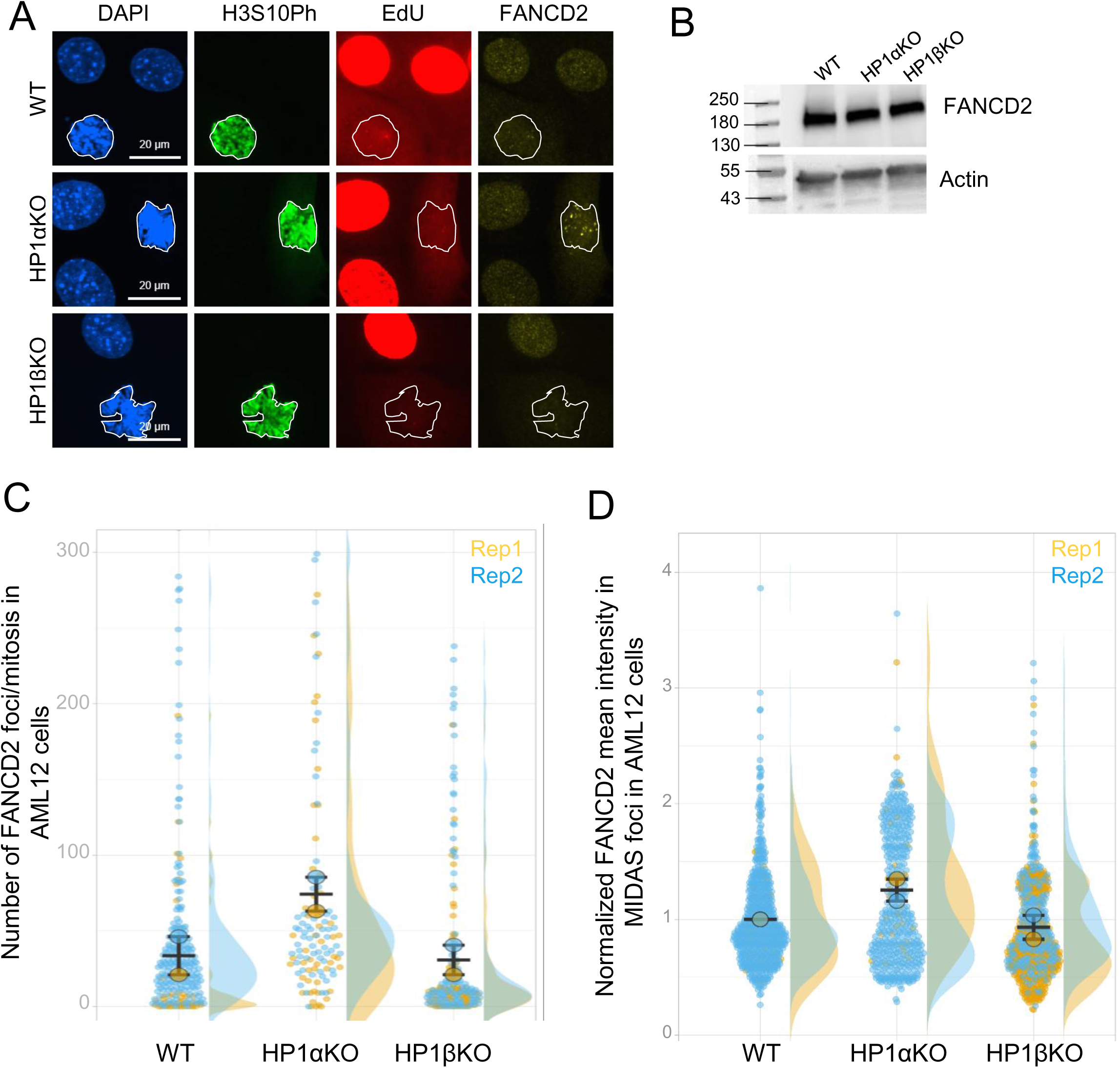
Impact of the loss of HP1α on FANCD2 expression and distribution towards MiDAS foci. (A) Representative image of IF analysis of FANCD2 expression in WT, HP1αKO and HP1βKO in interphasic (DAPI staining) and mitotic (DAPI and green staining, outlined in white) AML12 cells. EdU incorporation is in red (B) WB analysis of WT, HP1αKO and HP1βKO AML12 WCE using the indicated antibodies. (C) Superplot showing the distribution of the number of FANCD2 foci per WT, HP1αKO and HP1βKO AML12 mitotic nuclei as well as the mean number of FANCD2 foci per independent experiment. These data represent two independent experiments (rep1: WT, n=42, HP1αKO, n=52, HP1βKO, n=78 and rep2: WT, n=212, HP1αKO, n=68, HP1βKO, n=122). (D) Superplot showing the distribution of FANCD2 intensity per WT, HP1αKO and HP1βKO AML12 MiDAS foci per independent experiment as well as the normalized mean FANCD2 intensity to WT. These data represent two independent experiments (rep1: WT, n=22, HP1αKO, n=33, HP1βKO, n=380 and rep2: WT, n=641, HP1αKO, n=452, HP1βKO, n=268). Due to the limited number of independent experiments (n=2), no statistical analysis was performed for the number and the intensity of FANCD2 foci.

## Discussion

In this study, we demonstrated that some specific genomic regions display HP1- dependent instability in response to mild replicative stress induced by aphidicolin treatment. Interestingly, depending upon the localization of the APH-induced chromosomal breaks, their sensitivity towards HP1α and/or HP1β is different. Indeed, we found that the number of chromosomal breaks in response to replication stress is similarly affected by the loss of HP1α or HP1β around pericentromeric heterochromatin, whereas the number of breaks within chromosome arms, that most likely display euchromatic features is specifically affected by the loss of HP1α but not of HP1β, suggesting that the specific protective function of HP1α described here is at the level of euchromatic regions deprived of H3K9me3 whereas the protective functions shared between HP1α and HP1β is rather specific to H3K9me3-enriched constitutive heterochromatic regions. This conclusion is in line with the previously described HP1 functions in double strand break repair within heterochromatin and the demonstration that drosophila HP1a has H3K9-dependent and independent functions in the 3D organization of the genome (27,28). Furthermore, although chromosomal breaks within arms of HP1αKO metaphasis can be rescued by re-expression of exogenous HP1α protein, PCH breaks cannot be rescued in this same condition. Intriguingly, re-expression of a mutant HP1α (V22M) unable to interact with H3K9me3 is able to rescue the phenotype of chromosomal breaks both in the arms and at PCH of HP1αKO metaphasis. These results suggest that the chromatin environment surrounding HP1-dependent chromosomal breaks is essential for their stability however, the mechanistic basis of this unexpected observation remains to be determined.

Interestingly, the increased number of chromosomal breaks induced by APH observed in absence of HP1α correlates with an increased number of mitotic cells still undergoing DNA replication (MiDAS). Our data showed that this increased MiDAS activity in HP1αKO cells is likely associated with a slower replication fork progression and an increased recruitment of the Fanconi anaemia factor FANCD2 at these loci in absence of HP1α. These observations resemble those made for common fragile sites (CFS) (29,30). Therefore, our data suggest that the loss of HP1α allowed to uncover genomic regions that behave as HP1α-dependent CFS (13). These data indicate that, in addition to the previously recognized roles of HP1α and HP1β in DNA repair within heterochromatin, HP1α appears to have specific functions at some HP1α- dependent CFS most likely throughout the euchromatic compartment, independently of its ability to associate to H3K9me3. Interestingly these results are conserved between both the human HepG2 cells and the mouse AML12 cells, suggesting that this function towards these specific genomic loci is a hallmark of HP1α functions. However, this specific HP1α function is in apparent contradiction with those published by Charaka et al. (2020) (31), who reported that loss of HP1β in MEFs and human cells leads to increased γH2AX foci, replication fork stalling and firing of new origins of replication, as well as increased chromosomal gaps and breaks in response to hydroxyurea or cisplatin treatment. Although we also observed increased γH2AX foci in HP1βKO AML12 cells, several key differences between the two studies may explain the discrepancy in terms of chromosome alterations. First, the genotoxic agents used differ substantially in their mechanism of action and intensity since hydroxyurea induces a severe and global replication stress through nucleotide depletion and cisplatin generates direct DNA adducts, whereas we used a relatively low dose of aphidicolin that induces a selective replication stress, known to preferentially reveal naturally fragile genomic regions such as CFS. Second, Charaka et al. measured chromosomal gaps and breaks without distinguishing their chromosomal localization, whereas our analysis specifically separates breaks within pericentromeric heterochromatin from those occurring on chromosomal arms. It should however be noted that, in our study, gaps and breaks were not systematically distinguished and we cannot formally exclude that some gaps were included in our scoring. Taken together, these observations suggest that HP1α and HP1β may contribute to genome stability in response to severe genotoxic stress or direct DNA damage, consistent with their previously reported role in DNA repair within heterochromatin, whereas HP1α would have in addition, a specific and non-redundant protective function at discrete euchromatic loci that behave as CFS under mild replicative stress.

We further find that the recruitment of FANCD2, a factor known to be associated with CFS (26,29,30), is increased at MiDAS foci upon loss of HP1α. This result raises the possibility that HP1α and FANCD2 may compete for binding at foci that behave as HP1α-dependent CFS in mitosis. This conclusion is in line with what has been observed at sites of DNA damage. Indeed, Ayoub et al. showed that HP1β is first present at sites of DNA damage but has to be removed to allow the DNA damage response (32). Furthermore, the recruitment of HP1 proteins at sites of DNA damage has been shown to occur independently of H3K9me3, pointing to a H3K9me3-independent chromatin binding that may be a shared feature of HP1 functions in genome protection (33). Why HP1β is the main player in the DNA damage response whereas HP1α is specifically required for the protection against replication stress-induced breaks remains to be determined. What are the chromatin features that underly the protective role of HP1α at these specific genomic loci remains to be characterized and is beyond the scope of this study, but, as mentioned above, these regions are most likely deprived of H3K9me3. It is thus tempting to speculate that this specific function of HP1α is linked to its unique ability to form liquid-liquid phase separation (LLPS), whereas HP1β requires H3K9me3 to modified nucleosomes to fulfill this function (34–36).

## Materials and methods

### Cell culture

AML12 cells were cultured in DMEM+GlutaMAX (Gibco) supplemented with 10% fetal calf serum (FCS), 5 μg/ml insulin (Sigma), 5µg/ml human transferrin, 5ng/ml selenium and 40ng/ml Dexamethasone. HepG2 cells were cultured in DMEM+GlutaMAX (Gibco) supplemented with 10% fetal bovine serum. Cells were maintained at 37 °C and 5% CO2 in a humidified incubator. The absence of mycoplasma was verified every month in-house.

### Antibodies

The mouse anti-HP1, rabbit anti-TRIM28 and mouse anti-flag Abs were previously described and used at a 1:1000 dilution (37). Rabbit anti-γH2AX (Ab11174, Abcam; dilution 1:500), rabbit anti-53BP1 (Cell Signaling, 4937; dilution 1:500), rabbit anti-MCM10 (Antibodies, A88083, 1/500), rat anti-ORC2 (Santa Cruz, sc32734, 1/200); rat anti-ORC1 (Santa Cruz, sc23887, 1/200); rabbit anti-Chk1 phospho S345 (Cell Signaling, 2341, 1/1000), mouse anti-Chk1 (Cell Signaling, 2360, 1/1000); rabbit anti-FANCD2 (Ab108928, Abcam, dilution 1:400), rat anti- H3S10Ph (1:400); Rabbit anti-histone H3 (Abcam, 1731, 1/1000).

### Western blotting

Cells washed with phosphate-buffered saline (PBS) were lysed in 2X SDS Sample Buffer (10 mM Tris-HCl pH6.8, 5% glycerol, 1% SDS, 1 mM DTT, 0.5% β-mercapto-ethanol) at 95°C for 10 minutes. After measuring protein quantity by Bradford, equal amounts of protein were resolved by SDS-PAGE, transferred to a nitrocellulose membrane, and probed with the indicated antibodies. For fractionation, cells were resuspended with Buffer A (10 mM Hepes pH 7.9, 10 mM KCl, 1.5 mM MgCl2, 0.34 M Sucrose, 10% Glycerol, 1 mM DTT, 0.1% TritonX- 100, and protease inhibitors (Roche) and incubated for 2 minutes on ice to extract cytoplasmic soluble proteins. After centrifugation, pellets containing nuclei were resuspended with Buffer B (3 mM EDTA, 0.2 mM EGTA) and incubated for 30 minutes with occasional vortex on ice to extract nuclear soluble proteins. After centrifugation, supernatant containing nuclear soluble proteins was collected (S2) and cells were washed with the Buffer B. Then, chromatin (Chr) was dissolved in 2X SDS Sample Buffer (10 mM Tris-HCl pH6.8, 5% glycerol, 1% SDS, 1 mM DTT, 0.5% β-mercaptoethanol) and resolved by SDS-PAGE before immunoblotting. Secondary antibodies used are the following: Anti-rabbit 1:10000 (Cell Signaling Technology) and anti-mouse 1:10000 (Cell Signaling Technology) coupled to HRP. Protein bands were visualized on X-ray film by electro-chemiluminescence (Immobilon Western HRP Substrate, WBKLS0500, Millipore).

### CRISPR/Cas9-mediated HP1α and/or HP1β inactivated cellular models

Knock-out (KO) of *cbx5* (HP1α) and *cbx1* (HP1β) were generated by CRISPR-Cas9 technology in mouse HCC AML12 cell line and human HCC HepG2 cells. Briefly, the genome sequence corresponding to the mouse and human *cbx5* (HP1α) and *cbx1* (HP1β) genes were found in the NCBI website. The sgRNAs were designed using http://crispr.mit.edu site. sgRNAs were cloned at the BsmBI site in the lentiCRISPRv2 vector (Addgene) that harbours two cassettes for both expression of sgRNA and Cas9. After verification by sequencing, the lentiviral vector was used to produce viral particles in 293T cells. HepG2 and AML12 cells were infected and selected 48 h after infection in the presence of puromycin to obtain pools of HP1α- and/or HP1β depleted cells. Cells were then diluted and grown until individual clones could be picked and amplified for further characterization. The gRNA sequences are in the Supplementary table S1.

### Metaphase spreads

Cells were plated at 1.5.10^6^ cells/10 cm plates. The next day cells were treated with 1.2µM APH or the vehicle (3.6µl DMSO) for 20h and incubated 2 to 4h with 3µM Nocodazole (Sigma). Cells were harvested and resuspended in 10ml of pre-warmed hypotonic solution (14% FCS, 10.7mM KCl) and incubated 10 minutes at 37°C. 500µl of fixative solution (1 vol glacial acetic acid + 3 vol of 100% EtOH) were added to the hypotonic solution, centrifuged 5 minutes at 1000rpm and resuspended in 7ml of cold fixative solution. After centrifugation, the pellets were resuspended in 4ml of cold fixative solution and kept at least 12h at -20°C until use. 15µl of these cells were spread on cold superfrost slides from about 20cm above the slides, left to dry on a plate at 55°C for 2h and mounted with ProLong Diamond antifade containing DAPI (Invitrogen). Images were acquired with the Thunder microscope with 63X objective immersion oil (Leica) and the chromosome breaks for most experiments were counted twice, once by the experimenter and once by FC.

### DNA fibers

To label DNA, cells in exponential phase of growth were treated 30 minutes with 25μM 5-iodo- 20-deoxyuridine (IdU) (Sigma-Aldrich, 17125) and washed with medium, then treated with 50μM 5-chloro-20-deoxyuridine (CldU) (Sigma-Aldrich, C6891) for 30 minutes. After labelling, cells were collected with trypsin and resuspended in PBS at 500 cells/μl. Cells were lysed with spreading buffer (0.5% SDS, 200 mM Tris-HCl, pH 7.5, and 50 mM EDTA) and DNA fibers were stretched onto glass slides and left to air dry then fixed in methanol/acetic for 10 minutes. After fixation, DNA was denatured with 2.5 M HCl for 1h 30 minutes, and slides were blocked with 6%BSA PBS 0.1% Tween-20 for 1 h. Then, slides were incubated with primary anti-BrdU/IdU (BD Biosciences, 34780) and anti-CldU (Origene, TA190126) for 45 minutes (1/100) followed with secondary antibodies Alexa 488 anti-mouse (Invitrogen, A11017) and anti-rat- Cy3 (Jackson Immuno-research, 712-166153) respectively for 30 minutes (1/100) in PBS 0.1% TritonX-100 at 37°. Slides were mounted with ProLong Diamond antifade (Invitrogen). Fibers were visualized and imaged by Zeiss Axio Imager M2 upright microscope (Zeiss) equipped with an Apochromat 40X objective (NA 1.4, immersion oil). Replication track lengths were analyzed using ImageJ software. Statistical significance was performed using GraphPad Prism by a paired t-test on the median length of the tracks.

### Immunofluorescence and Microscopy analysis

5.10^5^ cells were seeded on glass coverslips in 1cm well plates. When required (γH2AX and 53BP1), cells were pre-extracted with ice-cold PBS containing 0.1% Triton-X 100 for 1-2 minutes and fixed with 4% formaldehyde (Thermo Scientific) diluted in PBS for 15 minutes at RT. After fixation, cells were permeabilized with PBS 0.25% Triton-X 100 for 10 minutes at RT. Then, cells were incubated in 5%BSA PBS 0.1% Tween20 for 1h at RT prior to incubation overnight at 4°C with the primary antibodies diluted in 5% BSA PBS 0.1% Tween20 ON at 4°C. On the following day, cells were washed with PBS 0.5% Tween-20 and incubated with appropriate secondary mouse and rabbit antibodies coupled to Alexa Fluor 568 or 488 (Thermo Fisher Scientific) respectively diluted in 5% BSA PBS 0.1% Tween20 (1/1000) for 2h at RT. DNA was counterstained with 0.1 µg/ml of DAPI (DNA intercalator, 4’,6-Diamidino-2-Phenylindole, D1306, Invitrogen). Coverslips were mounted with ProLong Diamond Antifade (Invitrogen). For imaging, images were acquired with an ORCA-Flash4.0 LT+ Digital CMOS camera (C1140-42U30, Hamamatsu) controlled by Zen acquisition software (Zeiss) using Zeiss Axio Imager M2 upright microscope (Zeiss) equipped with an Apochromat 63X objective (NA 1.4, immersion oil) or with thunder microscope (Leica). Images were prepared using Zen software and nuclear foci were quantified using CellProfiler^TM^ software. Representative images were prepared using Photoshop software (Adobe). Statistical significance was evaluated using GraphPad Prism by an unpaired t-test or two-tailed Student t-test on the median intensity of the labeling

### MiDAS

5.10^5^ cells were seeded on glass coverlids in 1cm well plates. The next day, cells were treated with 1.2µM APH and 4h later 10µM of the CDK1 inhibitor RO-3306 was added for further 16h. Cells were washed twice with fresh medium and treated with 10µM EdU for 30 minutes and fixed with 10% formalin (VWR). Cells were then first submitted to IF as described above and further submitted to the click-iT according to the Click-It ® EdU imaging kit manufacturer’s indications (Thermo Fisher Scientific). For imaging, images were acquired with the Thunder microscope with 63X objective immersion oil (Leica). Images analysis was made as follows: CellProfiler 4.2.8 was used to quantify immunofluorescence images. Metadata were extracted from the original .lif file names. Fluorescence channels were assigned accordingly. Nuclear segmentation was performed using the deep learning algorithm Cellpose 3.0.4 ((38) within CellProfiler (*RunCellpose* module, https://github.com/CellProfiler/CellProfiler-plugins). For mitotic cells, segmentation was carried out using classical intensity thresholding (Otsu), based on the phospho-H3(Ser10) staining. Nuclear MiDAS and/or FANCD2 foci were detected using the IdentifyPrimaryObjects module following an enhancement step (EnhanceOrSuppressFeatures, speckles, feature size = 10). Foci were segmented using a manual thresholding strategy and subsequently assigned to their corresponding nuclei using the *RelateObjects* module. Fluorescence intensities of the different channels were quantified within nuclei and/or individual foci using the *MeasureObjectIntensity* module. Segmentation quality and foci detection accuracy were systematically verified for every image using the *GrayToColor* and *OverlayOutlines* modules, allowing the reconstruction of the images and visual confirmation of the detected nuclei and foci.

Quantitative data were exported as .csv files using the ExportToSpreadsheet module and subsequently analyzed using GraphPad Prism 10 software. For comparisons involving more than two groups, statistical analyses were performed using ordinary one-way ANOVA or Kruskal-Wallis tests with Dunn’s post hoc correction, depending on data normality assessed using Shapiro–Wilk tests. Mann-Whitney tests were conducted for pairwise comparisons. Statistical significance was defined as follows: ns: not significant; *: p<0.05; **:p<0.01; ***:p<0.001; ****:p<0.0001. Image analysis was performed by QuantImagin (Tom EGGER, Montpellier, France.

## Supporting information

This article contains supporting information

## Author Contributions

K.Y. was involved in immunofluorescence experiments and FACS analyses, T.N.N. performed metaphase analysis and replication analysis, E.J. helped in critical reading and financial support, F.C. designed the study, obtained the main financial support, was involved in performing the MiDAS experiments and in the writing of the manuscript.

The authors used Claude (Anthropic) as an AI writing assistant to help structure and refine the language of this manuscript. All scientific content, interpretations and conclusions are the sole responsibility of the authors.

