## Supplementary Figure 1 for "Isotype specific loss of HP1α but not of HP1β uncovers genomic regions that behave as HP1α-dependent common fragile sites"

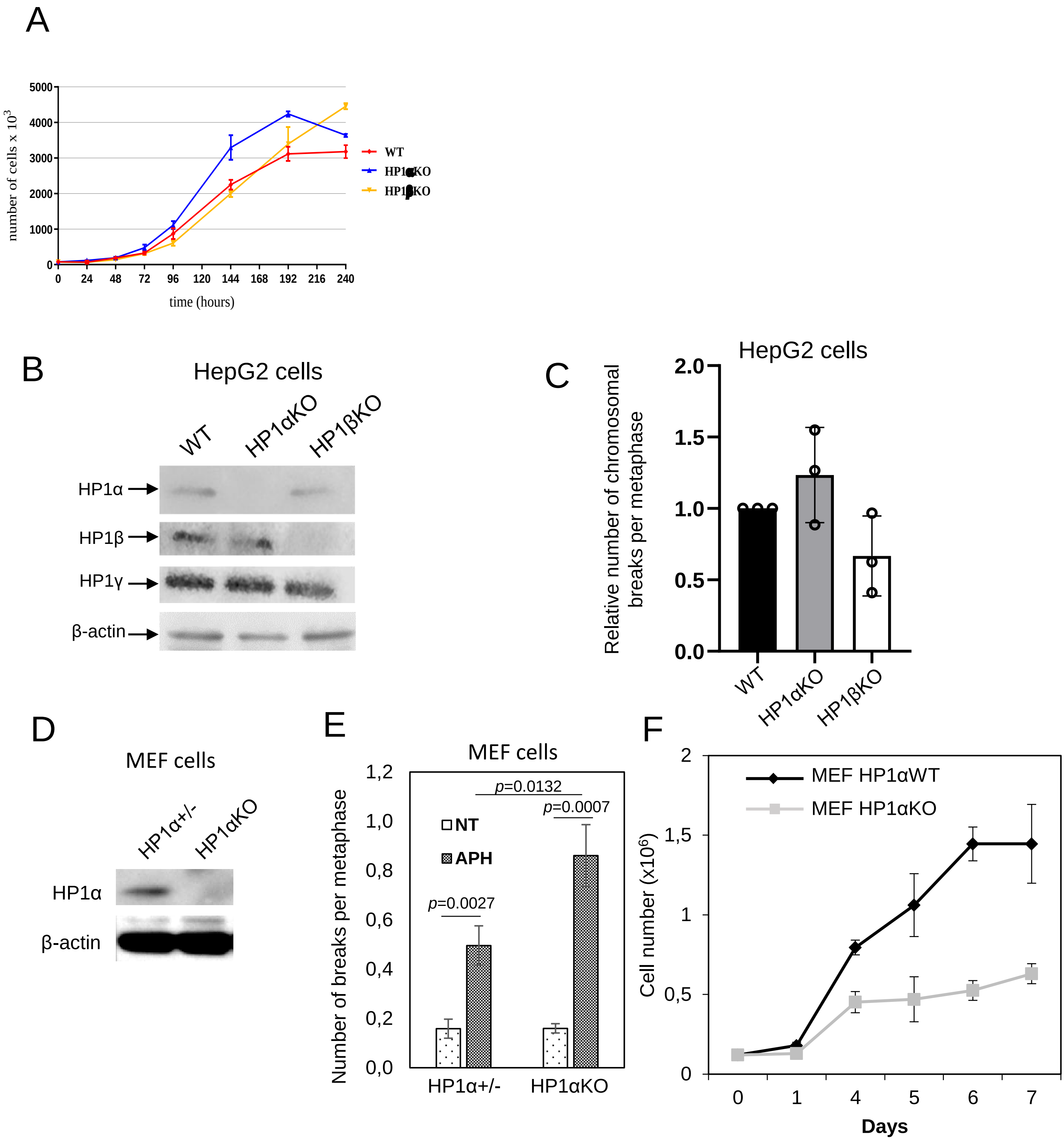


**Figure S1** : HP1α but not HP1β is involved in the stabilization of specific genomic loci in response to aphidicolin-induced replication stress. (A) Proliferation of AML12 WT, HP1αKO and HP1βKO cell lines in absence of APH treatment. (B) WB analysis of WCE from human Hepato Cellular Carcinoma HepG2 WT, HP1αKO and HP1βKO cell lines with the indicated antibodies (C) Quantification of the number of chromosomal breaks in HepG2 WT, HP1αKO and HP1βKO cell lines treated for 20h with 1.2µM APH. The mean values for each condition were normalized to WT in each independent experiment (n=3). An average of around 40 metaphases were counted per condition. Statistical significance was assessed using a one-sample t-test against the theoretical value of 1, the *p*-values correspond to comparison between WT and the different cell lines and were calculated using GraphPad Prism software. None of the comparison reached statistical significance. (D) WB analysis of WCE from Mouse Embryonic fibroblasts (MEF) *HP1*$\alpha^{+/-}$ and *HP1*αKO cell lines (E) Number of breaks per metaphase in three independent *HP1*$\alpha^{+/-}$ and *HP1*αKO MEF cell lines treated with 0.45µM APH. Statistical significance was calculated using student t-test (n=3). (F) Proliferation of WT and HP1αKO MEF cell lines in absence of APH treatment (n=2).
