## Supplementary Figure 2 for "Isotype specific loss of HP1α but not of HP1β uncovers genomic regions that behave as HP1α-dependent common fragile sites"

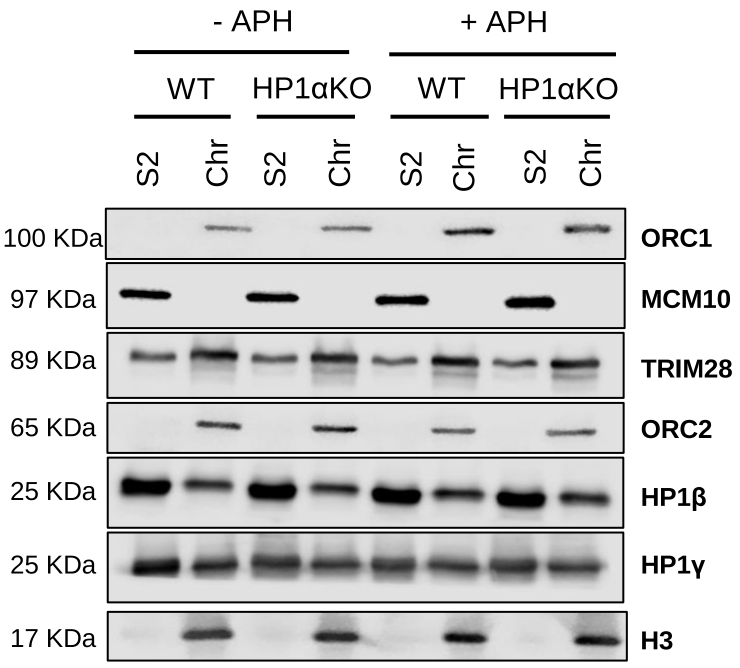


Figure S2: The level of expression and chromatin association of several proteins known to be involved in replication regulation and to interact with HP1 are not affected by the loss of HP1α in presence or absence of APH-induced replicative stress. WB analysis of soluble (S2) and chromatin (Chr) fractions of nuclear extracts from WT or HP1αKO AML12 treated or not with 1.2µM APH with the indicated antibodies
