## Supplementary Figure 3 for "Isotype specific loss of HP1α but not of HP1β uncovers genomic regions that behave as HP1α-dependent common fragile sites"

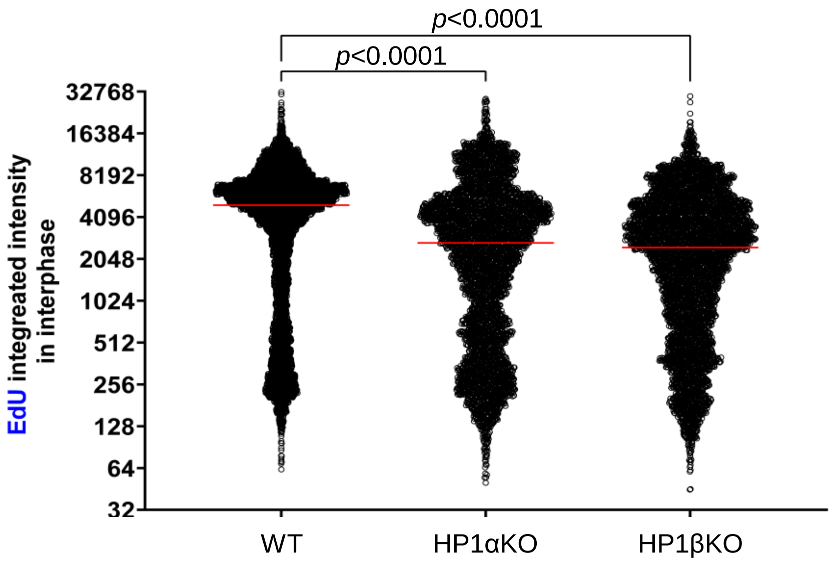


Figure S3: overall effect of loss of HP1α or HP1β on EdU incorporation in mouse hepatocyte AML12 interphasic cells. EdU intensity was quantified in WT, HP1αKO and HP1βKO AML12 interphasic nuclei using the CellProfiler software (see Materials and Methods). Statistical significance was calculated using Krustal-Wallis test, **p*<0,05; *****p*<0,0001
