## Supplementary Figure 4 for "Isotype specific loss of HP1α but not of HP1β uncovers genomic regions that behave as HP1α-dependent common fragile sites"

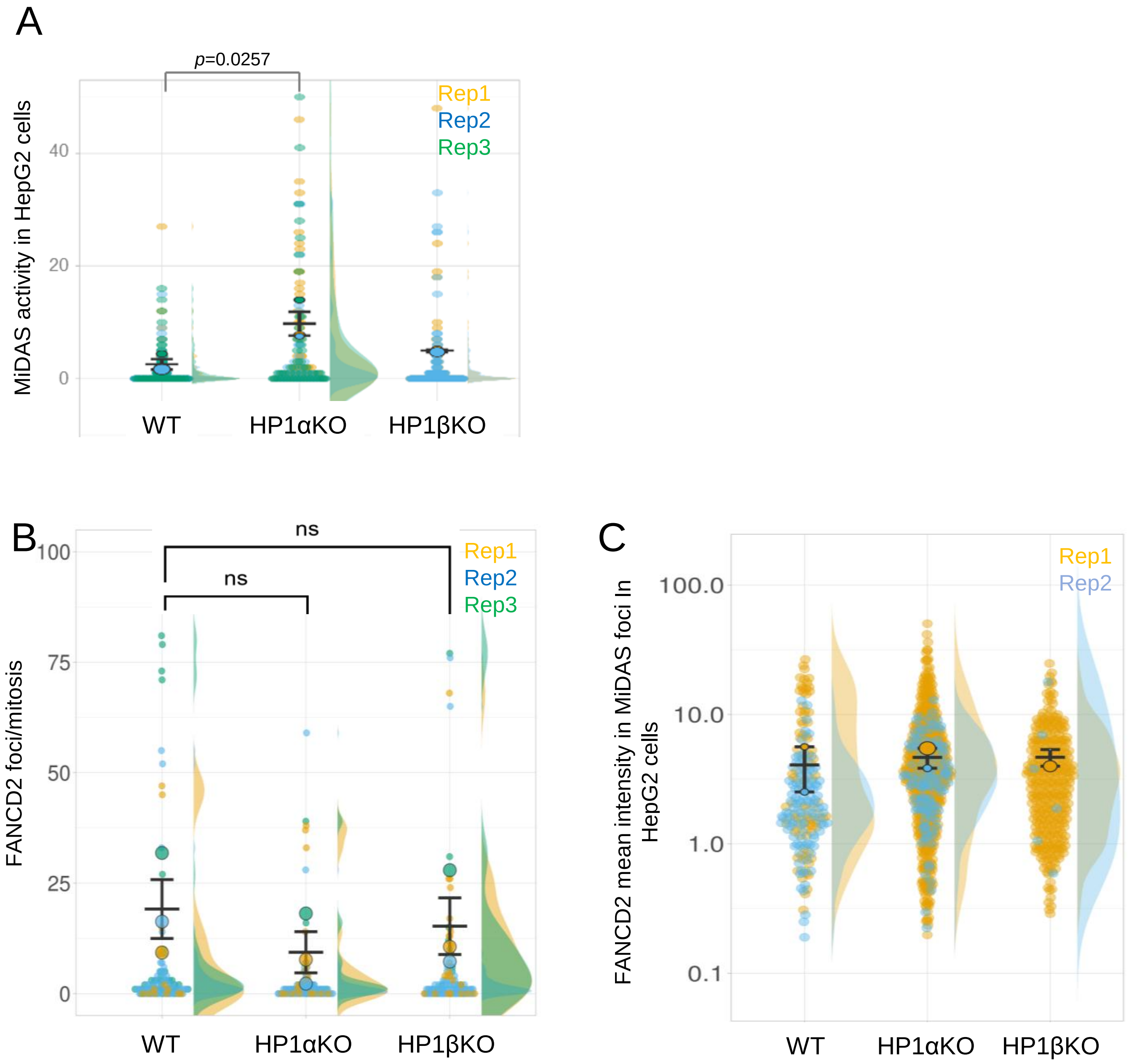


Supplementary figure S4 : FANCD2 analysis in HepG2 cells. (A) Superplot showing the distribution of the number of EdU foci as well as the mean number of EdU foci per WT and HP1αKO HepG2 mitotic nucleus for the three replicates (Rep1 in yellow, Rep2 in blue and Rep3 in green) and per HP1βKO mitotic nucleus for the two replicates (Rep1 in yellow, Rep2 in blue). Statistical analysis shown on the graph has been performed as paired t-test on the normalized means to WT of three independent experiments. (B) Superplot showing the distribution of FANCD2 intensity as well as the mean FANCD2 intensity per WT, HP1αKO and HP1βKO HepG2 mitotic nucleus for the three replicates (Rep1 in yellow, Rep2 in blue and Rep3 in green). Statistical analysis shown on the graph has been performed as Kruskal-Wallis test on data of three independent experiments. (C) Superplot showing the distribution of FANCD2 intensity as well as the mean FANCD2 intensity per MiDAS foci in WT, HP1αKO and HP1βKO HepG2 mitotic nuclei for the two replicates (Rep1 in yellow and Rep2 in blue). No statistical analysis was performed on these 2 independent experiments.
