## Supplementary table 1 for "Isotype specific loss of HP1α but not of HP1β uncovers genomic regions that behave as HP1α-dependent common fragile sites"

| target | primer name | sequence |
|---|---|---|
| mouse cbx5 | gRNA1-cbx5-m-fwd | CACCGTTGGACAGGCGCATGGTTAA |
| | gRNA1-cbx5-m-rev | AAACTTAACCATGCGCCTGTCCAAC |
| mouse cbx1 | gRNA1-cbx1-m-fwd | CACCGTGGTGGAAAAAGTTCTTGAT |
| | gRNA1-cbx1-m-rev | AAACCTGGAGTCAGTAGCTCCAATC |
| human cbx5 | gRNA-cbx5-h-fwd | CACCGCTAGACAGGCGCGTGGTTAA |
| | gRNA-cbx5-h-rev | AAACTTAACCACGCGCCTGTCTAGC |
| human cbx1 | gRNA-cbx1-h-fwd | CACCGAAAAGTTCTCGACCGTCGAG |
| | gRNA-cbx1-h-rev | AAACCTCGACGGTCGAGAACTTTTC |

Table S1 : sequences of gRNA used for for CRISPR-Cas9 inactivation of HP1α and HP1β-encoding genes
